# *Yersinia pseudotuberculosis* growth arrest during type-III secretion system expression is associated with altered ribosomal protein expression and decreased gentamicin susceptibility

**DOI:** 10.1101/2024.09.02.610769

**Authors:** Justin Greene, Rhett A. Snyder, Katherine L. Cotten, Ryan C. Huiszoon, Sangwook Chu, Rezia Era D. Braza, Ashley A. Chapin, Justin M. Stine, William E. Bentley, Reza Ghodssi, Kimberly M. Davis

## Abstract

It has been long appreciated that expression of the *Yersinia* type-III secretion system (T3SS) in culture is associated with growth arrest. Here we sought to understand whether this impacts expression of ribosomal protein genes, which were among the most highly abundant transcripts in exponential phase *Yersinia pseudotuberculosis* based on RNA-seq analysis. To visualize changes in ribosomal protein expression, we generated a fluorescent transcriptional reporter with the promoter upstream of *rpsJ*/S10 fused to a destabilized *gfp* variant. We confirmed reporter expression significantly increases in exponential phase and decreases as cells transition to stationary phase. We then utilized a mouse model of systemic *Y. pseudotuberculosis* infection to compare T3SS and S10 reporter expression during clustered bacterial growth in the spleen, and found that cells expressing high levels of the T3SS had decreased S10 levels, while cells with lower T3SS expression retained higher S10 expression. In bacteriological media, growth inhibition with T3SS induction and a reduction in S10 expression were observed in subsets of cells, while cells with high expression of both T3SS and S10 were also observed. Loss of T3SS genes resulted in rescued growth and heightened S10 expression. To understand if clustered growth impacted bacterial gene expression, we utilized droplet-based microfluidics to encapsulate bacteria in spherical agarose droplets, and also observed growth inhibition with high expression of T3SS and reduced S10 levels that better mirrored phenotypes observed in the mouse spleen. Finally, we show that T3SS expression is sufficient to promote tolerance to the ribosome-targeting antibiotic, gentamicin. Collectively, these data indicate that the growth arrest associated with T3SS induction leads to decreased expression of ribosomal protein genes, and this results in reduced antibiotic susceptibility.

**Author Summary:** Slow-growing bacterial cells have reduced antibiotic susceptibility, rendering them very difficult to eliminate during antibiotic treatment. However, for many key virulence factors (bacterial factors required to promote infection), it remains unclear whether expression is sufficient to slow bacterial growth and impact antibiotic susceptibility. Using *Yersinia pseudotuberculosis*, we found ribosomal protein expression fluctuated based on growth rate, and we generated a fluorescent reporter construct to detect altered ribosomal protein expression within individual bacterial cells. We then asked if expression of a key virulence factor in *Yersinia*, the type-III secretion system (T3SS), is sufficient to lower ribosomal protein expression, since it has been well established that T3SS induction results in growth arrest. We found high levels of T3SS expression promotes slowed growth and antibiotic tolerance, and bacterial cells that survive treatment with a ribosome-targeting antibiotic, gentamicin, have heightened levels of T3SS and lower levels of S10 expression.

## Introduction

Exposure to host-derived antimicrobials and immune cell interaction triggers expression of bacterial virulence factors, components required to combat host defenses and promote disease. While virulence factors are necessary during infection, high levels of expression can alter bacterial cell physiology, and impact overall fitness to the point of promoting slowed growth and reduced antibiotic susceptibility (1). Further, virulence factor expression is often heterogeneous across a bacterial population which may promote differential antibiotic susceptibility amongst individual cells (1–3). Mixed populations of rapidly-growing, antibiotic-susceptible cells and slowly-growing, antibiotic-tolerant cells are termed antibiotic persistent populations (4, 5). It can be very difficult to fully eliminate these populations during antibiotic treatment, resulting in residual subpopulations of bacterial cells capable of causing relapsing infection. It has also been shown that persistent populations can give rise to antibiotic resistant cells (6, 7), linked to prolonged exposure of persistent cells to host-derived stressors and antibiotic stress during courses of antibiotic treatment. For these reasons, it is critical to better understand the mechanisms underlying alterations in antibiotic susceptibility and explore treatment regiments which are more effective at eliminating the entirety of bacterial populations.

One of the central virulence factors for Gram-negative bacterial pathogens is a type-III secretion system (T3SS), which injects its associated effector proteins into the host cell cytoplasm to modulate host cell processes (8). For human pathogenic *Yersiniae* (*Yersinia pseudotuberculosis, Yersinia enterocolitica, Yersinia pestis*), the T3SS effectors uncouple host signaling cascades, limit reactive oxygen species production, and prevent phagocytosis to allow bacteria to remain in an extracellular niche (9–13). Expression of the *Yersinia* T3SS has been shown to arrest growth in culture (14, 15); however, it remains unclear if expression is linked to slowed growth in mouse models of infection, or if T3SS expression is sufficient to reduce antibiotic susceptibility. These questions have been difficult to assess with bacterial genetic approaches, because of limited host immune cell infiltration in the absence of T3SS (16, 17), which simultaneously removes potential fitness costs and selective pressures. Moreover, the link between T3SS expression and antibiotic susceptibility has only been explored in the context of *Salmonella enterica* serovar Typhimurium (*S.* Typhimurium) SPI-1, a T3SS which, in contrast with *Yersinia*, promotes bacterial invasion (18). Thus, it remains unclear if expression of T3SSs more broadly impact the antibiotic susceptibility of Gram-negative pathogens.

*Y. pseudotuberculosis* is a natural pathogen of rodents and humans, and has been used since the 1960’s to study different aspects of host-pathogen interactions, such as host cell invasion (19), dynamics of virulence factor expression (20–22), dissemination and bacterial population bottlenecks (23–25), generation of immune memory in gastrointestinal-associated lymphoid tissues (26, 27), inflammasome activation (16, 17), and several different aspects of T3SS biology, including structure determination (28, 29), host cell targeting (30, 31), and regulation of expression (22, 32, 33). Knowledge gained from these studies has provided critical insight into host-pathogen interactions. In the mouse model of *Y. pseudotuberculosis* systemic infection, we have previously shown that extracellular bacteria replicate to form clonal clusters, called microcolonies, in the spleen (20). Bacteria at the periphery of microcolonies are in direct contact with neutrophils and express very high levels of the T3SS (20). Bacteria at the center of microcolonies lack host cell contact and have an intermediate level of T3SS expression, which is induced at mammalian body temperatures (37° C) (22, 34). In addition to expressing high levels of T3SS, bacteria at the periphery of microcolonies are also exposed to other host-derived stressors, such as high levels of nitric oxide (NO), which is sufficient to slow the growth of bacteria and impact antibiotic susceptibility. In previous studies, we identified peripheral cells exposed to NO and showed they exhibited slowed growth and decreased antibiotic susceptibility (3, 21, 35). However, it remained unclear whether heightened T3SS expression contributed to these phenotypes alongside NO exposure.

Here, we first characterized the transcriptional signature of exponential phase *Y. pseudotuberculosis* and detected high levels of ribosomal protein transcripts in rapidly growing cells. We then generated a transcriptional reporter by fusing the promoter upstream of ribosomal protein operon *rpsJ*/S10 to destabilized *gfp* and showed that rapidly growing cells have heightened S10 reporter expression. Within the mouse spleen, cells with lower levels of T3SS expression exhibited higher levels of S10, while cells with higher levels of T3SS expression had lower S10 levels in the spleen. We then used two systems to model differential T3SS expression in culture: planktonic growth in bacteriological media and an agarose droplet system, which models clustered bacterial growth. We show that high levels of T3SS expression are sufficient to slow bacterial growth, reduce S10 expression levels, and reduce susceptibility to the ribosome-targeting antibiotic, gentamicin.

## Results

### Exponential phase cells express high levels of ribosomal protein genes

We first utilized RNA-seq to determine the transcriptional profile of exponential phase *Yersinia pseudotuberculosis* as compared to stationary phase cells. This approach would allow us to identify a promoter region that would be specifically activated in rapidly growing, exponential phase cells. Wild-type (WT) *Y. pseudotuberculosis* cells were cultured overnight (stationary phase) and subcultured in fresh media (exponential phase) at 26° C to avoid spontaneous virulence plasmid loss, which can occur at a relatively high frequency during prolonged culture at 37° C (36–38). We found 894 transcripts increased more than 2-fold in exponential phase, 806 genes increased more than 2-fold in stationary phase, and 2255 protein coding genes were unchanged (**S1 Table, S2 Table**). KEGG pathway analyses indicated that transcripts for genes associated with the ribosome, biosynthesis of secondary metabolites, and aminoacyl-tRNA biosynthesis were heightened in exponential phase (**Fig 1A**), while transcripts for flagellar assembly and two-component systems were heightened in stationary phase (**Fig 1B**). Additional groups of genes expressed in stationary phase included genes involved in tyrosine metabolism, ABC transporters, degradation of aromatic compounds, and quorum sensing (**Fig 1B**). Enrichment of these KEGG pathways was somewhat expected, based on RNA-seq analyses in other *Y. pseudotuberculosis* strains (39). Of note, we found transcript levels of 56 genes were greater than 10-fold more abundant in exponential compared to stationary phase, and 22 of these 56 genes were ribosomal protein genes (**Table 1**). This set included *rpsJ*, which encodes for the 30S ribosomal protein S10, and whose transcripts were ∼13 fold more abundant in exponential phase. *rpsJ* is the first gene expressed in a large operon of small and large ribosomal subunit protein genes (**Fig 1C**), suggesting that activity of the promoter upstream of *rpsJ* could serve as a reporter for transcription of the operon. Consistent with this assumption, all genes in this operon (encoding for: S10, L3, L4, L23, L2, S19, L22, S3, L16, L29, S17) were among the top 56 most highly upregulated genes in exponential phase, with more than 10-fold increased transcript abundance in exponential phase cells (**Fig 1C**, **Table 1**). Additionally, *rpsJ*/S10 expression is transcriptionally regulated by DksA and ppGpp in stationary phase, indicating a reporter containing the *rpsJ* leader and promoter sequences could reflect DksA and ppGpp-dependent transcriptional regulation of other ribosomal genes as well (40). For these reasons, we chose the leader and promoter sequences upstream of the S10 ribosomal protein gene (*rpsJ*) for our ribosomal reporter construct to mark rapidly growing cells.

**Fig 1.**
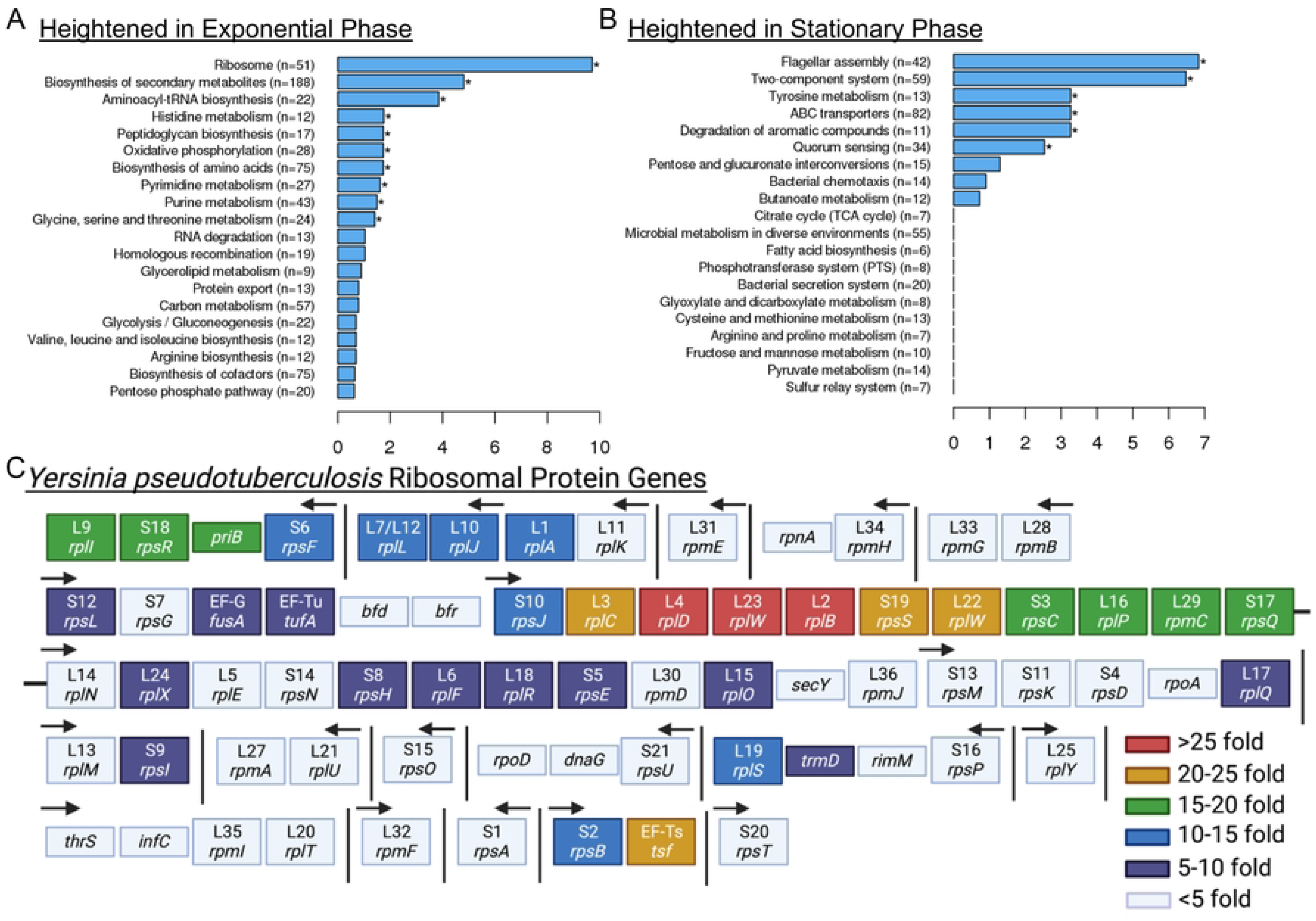
Exponential phase cells express high levels of ribosomal protein genes. RNA-seq was performed on exponential and stationary phase WT *Y. pseudotuberculosis* cells. The DESeq2 method of pairwise comparisons was used to determine significant differences in transcript levels. KEGG pathway analyses were utilized to determine the biological pathways overrepresented in **(A)** exponential and **(B)** stationary phase cells. **(C)** *Yersinia pseudotuberculosis* ribosomal protein gene organization. Protein subunit names and gene names are shown. Arrows indicate promoter regions, direction indicates direction of transcription, determined using genomes NZ_CP009712.1 and NZ_CP032566.1. Vertical lines indicate genes are not adjacent. Horizontal black line indicates S17/*rpsQ* and L14/*rplN* are adjacent. Genes are color-coded based on fold increase (>5) in exponential phase cells. Created in Biorender.

**Table 1:** Genes with highest fold change in exponential compared to stationary phase.

| log2foldchange | Fold change | padjust | gene_name | gene_description |
| --- | --- | --- | --- | --- |
| 5.696331523 | 51.8521363 | 1.34E-128 | ompC | porin OmpC |
| 5.352923512 | 40.8686735 | 4.86E-44 | DN756_RS01950 | hypothetical protein |
| 5.223072614 | 37.3509392 | 4.35E-44 | DN756_RS19535 | heme anaerobic degradation radical SAM methyltransferase ChuW/HutW |
| 4.90850355 | 30.0335592 | 1.46E-46 | rplD | 50S ribosomal protein L4 |
| 4.842504065 | 28.6905569 | 9.16E-50 | rplW | 50S ribosomal protein L23 |
| 4.747050226 | 26.8537233 | 1.68E-54 | rnpB | RNase P RNA component class A |
| 4.746679269 | 26.8468193 | 3.84E-54 | rplB | 50S ribosomal protein L2 |
| 4.699349382 | 25.9803576 | 1.33E-28 | hutX | heme utilization cystosolic carrier protein HutX |
| 4.624182865 | 24.6614012 | 1.22E-67 | rplV | 50S ribosomal protein L22 |
| 4.587681967 | 24.0452824 | 3.91E-18 | DN756_RS14930 | tRNA-Gln |
| 4.508151871 | 22.7556339 | 5.79E-49 | rpsS | 30S ribosomal protein S19 |
| 4.478526293 | 22.2931147 | 1.06E-18 | DN756_RS14935 | tRNA-Gln |
| 4.455238008 | 21.9361435 | 1.66E-79 | ilvC | ketol-acid reductoisomerase |
| 4.435700792 | 21.641083 | 6.14E-31 | tsf | elongation factor Ts |
| 4.402357007 | 21.1466468 | 1.53E-35 | rplC | 50S ribosomal protein L3 |
| 4.347487802 | 20.3574902 | 2.30E-67 | DN756_RS12480 | APC family permease |
| 4.302271957 | 19.729356 | 1.97E-17 | malK | maltose/maltodextrin ABC transporter ATP-binding protein MalK |
| 4.262147054 | 19.1881944 | 1.52E-101 | DN756_RS06760 | dihydrodipicolinate synthase family protein |
| 4.224799241 | 18.6978339 | 1.56E-124 | DN756_RS06765 | N-acetylmannosamine-6-phosphate 2-epimerase |
| 4.19204945 | 18.2781664 | 2.51E-110 | rpmC | 50S ribosomal protein L29 |
| 4.191842469 | 18.2755443 | 1.09E-53 | rpsR | 30S ribosomal protein S18 |
| 4.124259445 | 17.4391697 | 1.37E-91 | DN756_RS07545 | PTS sugar transporter subunit IIB |
| 4.058994111 | 16.6678268 | 7.52E-99 | rpsQ | 30S ribosomal protein S17 |
| 4.022704701 | 16.253795 | 4.77E-99 | rplI | 50S ribosomal protein L9 |
| 4.01661034 | 16.1852791 | 3.03E-87 | ompX | outer membrane protein OmpX |
| 4.006945136 | 16.0772097 | 8.21E-69 | priB | primosomal replication protein N |
| 3.999533604 | 15.9948283 | 7.87E-87 | rpsC | 30S ribosomal protein S3 |
| 3.939201347 | 15.3397317 | 1.44E-112 | rplP | 50S ribosomal protein L16 |
| 3.891815541 | 14.8440776 | 2.91E-20 | rpsB | 30S ribosomal protein S2 |
| 3.814396217 | 14.0684962 | 3.96E-22 | speA | biosynthetic arginine decarboxylase |
| 3.783491797 | 13.7703354 | 2.87E-77 | rplJ | 50S ribosomal protein L10 |
| 3.781213946 | 13.7486108 | 1.10E-107 | DN756_RS06775 | N-acetylmannosamine kinase |
| 3.738173255 | 13.3444992 | 1.75E-37 | htpG | molecular chaperone HtpG |
| 3.731351524 | 13.2815491 | 3.40E-28 | yidC | membrane protein insertase YidC |
| 3.721580822 | 13.1919033 | 2.49E-17 | DN756_RS06965 | tRNA-Val |
| 3.691318931 | 12.9180726 | 5.08E-30 | rpsJ | 30S ribosomal protein S10 |
| 3.647140184 | 12.5284861 | 1.80E-24 | DN756_RS19545 | SDR family oxidoreductase |
| 3.637044276 | 12.4411184 | 7.02E-17 | accC | acetyl-CoA carboxylase biotin carboxylase subunit carboxylase C-terminal domain |
| 3.625644565 | 12.3432 | 1.14E-67 | dnaJ | molecular chaperone DnaJ |
| 3.619695655 | 12.292408 | 1.03E-57 | rpsF | 30S ribosomal protein S6 |
| 3.605060389 | 12.1683394 | 4.38E-32 | hslV | ATP-dependent protease subunit HslV |
| 3.584315091 | 11.9946162 | 3.46E-184 | rplL | 50S ribosomal protein L7/L12 |
| 3.577533236 | 11.9383639 | 3.53E-123 | tig | trigger factor |
| 3.57208558 | 11.8933694 | 3.48E-19 | lpdA | dihydrolipoyl dehydrogenase |
| 3.563841838 | 11.8256029 | 5.08E-66 | rplS | 50S ribosomal protein L19 |
| 3.503267432 | 11.339361 | 3.51E-61 | DN756_RS12195 | TonB-dependent receptor |
| 3.494065924 | 11.2672687 | 3.55E-14 | DN756_RS06970 | tRNA-Val |
| 3.484338798 | 11.1915565 | 1.96E-29 | malP | maltodextrin phosphorylase |
| 3.461945036 | 11.0191806 | 9.74E-30 | DN756_RS19555 | TonB-dependent hemoglobin/transferrin/lactoferrin family receptor |
| 3.436664222 | 10.8277698 | 6.09E-93 | rplA | 50S ribosomal protein L1 |
| 3.383401419 | 10.435309 | 7.11E-138 | rplK | 50S ribosomal protein L11 |
| 3.366783823 | 10.3158001 | 2.82E-35 | yiuA | iron-siderophore ABC transporter substrate-binding protein YiuA |
| 3.357159614 | 10.2472125 | 9.38E-24 | cysK | cysteine synthase A |
| 3.355441448 | 10.235016 | 1.17E-23 | pyrH | UMP kinase |
| 3.35303223 | 10.2179383 | 2.04E-09 | cobA | uroporphyrinogen-III C-methyltransferase |
| 3.325103341 | 10.0220334 | 1.13E-25 | DN756_RS14130 | 2Fe-2S ferredoxin-like protein |

\*Colors correspond with highlighted fold change values in Fig 1C. *rpsJ*/S10 is outlined in black.

### Ribosomal reporter expression is heightened in exponential phase cells

To visualize S10 (*rpsJ*) expression dynamics throughout the cell cycle, an ectopic reporter construct was made using the S10 leader and promoter sequences to drive expression of a ssrA-tagged, destabilized GFP (21, 41, 42). The half-life of this destabilized version of GFP is approximately 59 minutes which will allow for visualization of dynamic changes in reporter expression (21). It is important to note this reporter was generated to detect rapidly growing exponential phase cells based on S10 ribosomal protein expression, in contrast with other approaches that detect ribosomal RNA, which may better mirror ribosome abundance (43, 44). After confirming ectopic GFP reporter expression did not affect growth rate (**Fig 2A**), *P_S10_::gfp-ssrA* cell populations were sampled at defined timepoints to represent different growth stages, and GFP fluorescent intensity within individual bacterial cells was quantified by fluorescence microscopy (**Fig 2B**). GFP expression significantly increased as cells transitioned from stationary phase (0h) to exponential phase (4h), and reporter signal significantly decreased at 8h (**Fig 2B**) as cells began to transition back into stationary phase. These results suggest that the S10 reporter can dynamically detect rapidly growing cells.

**Fig 2.**
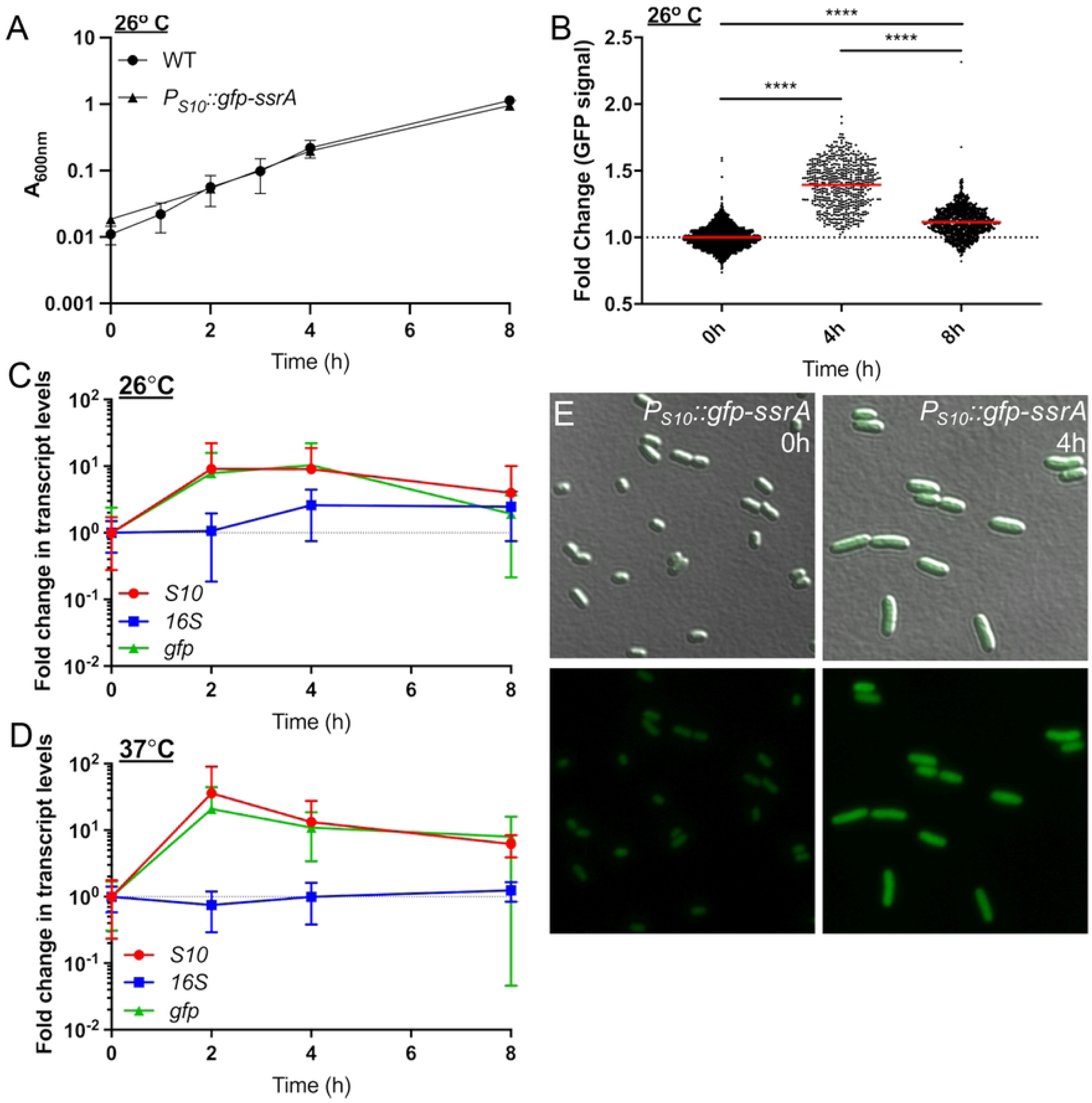
Ribosomal reporter expression is heightened in exponential phase cells. **(A)** Growth curve of WT and *P_S10_::gfp-ssrA Y. pseudotuberculosis* strains at 26° C. Absorbance (A_600nm_) detected by plate reader at the indicated timepoints (hours, h). Four biological replicates shown with mean and standard deviation. **(B)** Single cell fluorescence microscopy used to quantify the fold change in fluorescence of the *P_S10_::gfp-ssrA* strain at the indicated timepoints, sampled from cultures in **(A)**. Fold change in signal is relative to the mean GFP fluorescence value at 0h, which is shown as a dotted line at a value of 1. Horizontal lines: median values. Data represents 3 biological replicates for each condition. **(C-D)** qRT-PCR to detect *S10, gfp,* and *16S* transcript levels during culture of the *P_S10_::gfp-ssrA* strain at **(C)** 26° C or **(D)** 37° C. RNA was isolated at the indicated timepoints. Data represents 6 biological replicates. **(E)** Representative images used to quantify single-cell fluorescence corresponding with 0h and 4h timepoints in panel B. Statistics: **(B)** Kruskal-Wallis one-way ANOVA with Dunn’s post-test, ****p<.0001.

To confirm changes in GFP fluorescence reflect changes in S10 ribosomal protein transcript levels, qRT-PCR was performed to quantify transcript levels of *S10* and *gfp,* compared to ribosomal RNA (16S). Transcripts of the housekeeping gene, *rpoC*, were used here for normalization to detect any 16S fluctuation. *P_S10_::gfp-ssrA* cells were cultured overnight, sampled in stationary phase (0h), sub-cultured, and samples were collected at 2, 4, and 8 hours for RNA isolation. As mentioned above, 26° C is typically used for *Y. pseudotuberculosis* culture, since incubation of *Yersinia* at mammalian body temperatures (37° C) triggers expression of mammalian cell-targeting virulence factors, and prolonged culture can result in loss or inactivation of virulence genes (15, 36). To determine whether the S10 reporter would also detect faster-growing exponential phase cells during culture at 37° C, cells were grown in parallel at 26° C and 37° C. At 26° C, *gfp* and *S10* transcript levels increased between 0 and 2 hours, remained steady at 4h, and began to decline at 8h, while 16S transcripts remained relatively constant (**Fig 2D**). At 37° C, while 16S transcript levels stayed constant, both *gfp* and *S10* transcript levels peaked at 2h then declined between 4h and 8h, consistent with when bacteria were growing more rapidly (**Fig 2E**). *gfp* and *S10* transcript levels correlated well at each temperature and throughout the sampled timepoints, suggesting the *P_S10_::gfp-ssrA* reporter is accurately depicting *S10* transcript levels. Overall, these results indicate the S10 reporter can be utilized to detect rapidly growing exponential phase cells at either temperature.

### T3SS-high cells at the periphery of microcolonies have decreased ribosomal protein expression based on lower S10 reporter expression

We then sought to utilize the S10 reporter to determine if expression of critical virulence factors, specifically the type-III secretion system (T3SS) impacts the translational activity of individual cells based on S10 reporter expression. In the mouse model of systemic infection, we previously showed that *Y. pseudotuberculosis* at the periphery of microcolonies are in direct contact with neutrophils and express high levels of the T3SS (20). Bacteria at the center of microcolonies lack host cell contact, and have an intermediate level of T3SS expression, induced by growth at mammalian body temperatures (37° C) (22, 34). We hypothesized that the high levels of T3SS expression within peripheral cells would impact their fitness (14), potentially through a reduction of their total metabolic activity and decreased ribosomal content, as measured by S10 reporter expression.

To test this hypothesis, the S10 reporter construct was moved into a strain containing a fluorescent *yopE::mCherry* reporter to detect T3SS expression (20, 24). C57BL/6 mice were infected intravenously with the *yopE::mCherry P_S10_::gfp-ssrA* strain. Spleens were harvested at Day 2 and Day 3 post-inoculation to quantify bacterial colony-forming units (CFUs) and visualize reporter expression by fluorescence microscopy. Based on previous studies, we expected significant bacterial growth between these timepoints, allowing us to differentiate between slower-growing and faster-growing individual cells (21, 35). Bacterial CFUs slightly increased between these two timepoints, although this was not significant **(Fig 3A)**. The areas of bacterial microcolonies did not significantly increase between the timepoints, although more microcolonies were detected in the same number of cryosections of tissue at day 3 (2 cryosections/mouse), consistent with a low level of bacterial growth between the timepoints **(Fig 3B)**.

**Fig 3.**
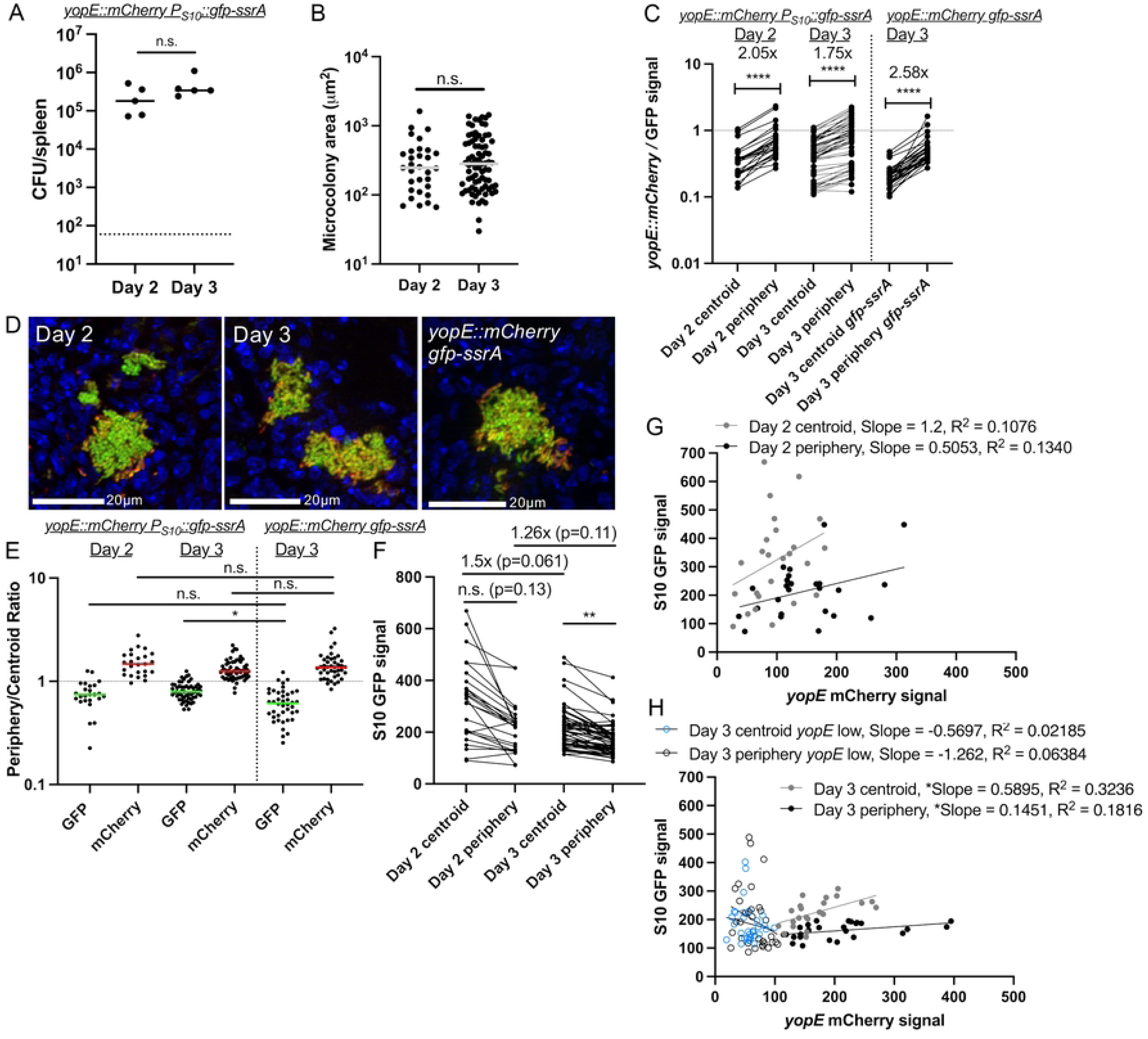
T3SS-high cells at the periphery of microcolonies have decreased ribosomal activity based on lower S10 reporter expression. C57BL/6 mice were inoculated intravenously with either the *yopE::mCherry P_S10_::gfp-ssrA* or *yopE::mCherry gfp-ssrA* (constitutive GFP) strains. **(A)** CFU/spleen was quantified for the *yopE::mCherry P_S10_::gfp-ssrA* strain at days 2 and 3 post-inoculation. Dots: individual mice. **(B)** Microcolony areas from spleens were quantified by fluorescence microscopy at the indicated timepoints. Dots: individual microcolonies. **(C)** Reporter expression (ratio mCherry/GFP signal intensity) was quantified at the indicated timepoints and spatial locations (centroid, periphery). Data with the S10 reporter strain is shown on the left with constitutive GFP on the right. Horizontal dotted line represents a value of 1, equal amounts of mCherry and GFP signal. **(D)** Representative images of microcolonies: day 2 and day 3 *yopE::mCherry P_S10_::gfp-ssrA* shown alongside day 3 *yopE::mCherry gfp-ssrA.* **(E)** Microcolony reporter expression quantified by the periphery/centroid ratio of either GFP or mCherry signal. **(F)** S10 GFP signal quantified for microcolonies of the *yopE::mCherry P_S10_::gfp-ssrA* strain at the indicated timepoints and spatial locations. **(G, H)** Linear regression data indicating slope and R2 value of best fit lines for GFP compared to mCherry signal of microcolonies at the indicated spatial locations at **(G)** day 2 and **(H)** day 3. **(H)** Microcolonies with low levels of *yopE*::mCherry signal were analyzed separately (open circles). Dots represent individual mice **(A)** or individual microcolonies (all other panels). Statistics: **(A-B)** Mann-Whitney; **(C)** Wilcoxon matched-pairs; **(E, F)** Kruskal Wallis one-way ANOVA with Dunn’s post-test; ****p<0.0001, **p<.01, *p<.05, ns: not-significant. **(G, H)** Significantly non-zero slope indicates correlation between values.

To analyze S10 reporter expression, we performed infections in parallel with a control strain, *yopE::mCherry gfp-ssrA* that constitutively expresses unstable GFP. This control is particularly important within mouse tissues because of the spatial architecture of microcolonies; fluorescent signals at the centroid can appear higher due to increased bacterial cell density at this location relative to the periphery. Because we are expecting this pattern based on our hypothesis, we wanted to ensure that any observed patterning was truly due to changes in reporter expression and not a result of cell density differences, and so we analyzed microcolonies from mice infected with the control strain (*yopE::mCherry gfp-ssrA*) at day 3 compared to *yopE::mCherry P_S10_::gfp-ssrA* microcolonies. Following imaging of tissues, a ratio of the mCherry (*yopE*) to GFP fluorescence was quantified for each strain at each microcolony centroid and periphery. For all microcolonies analyzed, the mCherry/GFP ratio was heightened at the periphery relative to the centroid **(Fig 3C)**. The spatial differences in the mean mCherry/GFP ratios (centroid vs. periphery) were most pronounced with the control strain (2.58x increase) and were lower with the S10 reporter strains (2.05x at day 2, 1.75x at day 3) **(Fig 3C)**. This indicates S10 (*rpsJ*) expression differs from constitutive expression, and is instead spatially-regulated across microcolonies. Interestingly, the mCherry/GFP ratios between the centroid and periphery become more similar at day 3 when compared to day 2 **(Fig 3C, 3D)**. This could indicate S10 expression is decreasing at the centroid by day 3, reflecting slowed growth, or that the periphery was in fact retaining ribosomal protein expression.

To investigate this temporal shift in mCherry/GFP ratio between the centroid and periphery of microcolonies, we quantified the ratio of periphery-to-centroid signal for each fluorescent channel independently. As expected (20), *yopE* reporter values were consistently above a value of 1, indicating similar levels of peripheral expression across both strains and timepoints. **(Fig 3E**). In contrast, S10 values were primarily below a value of 1, indicating more centroid expression. While GFP fluorescence signals were similar between constitutive expression and the S10 reporter at day 2, the S10 reporter GFP values were significantly higher than constitutive GFP at day 3 **(Fig 3E**). This ratio change could reflect that S10 reporter expression is increased at the periphery on day 3, or that S10 levels are decreasing at the centroid, consistent with slowed growth.

We then analyzed S10 reporter expression independently between the centroid and periphery of microcolonies and found the S10 mean reporter signal was 1.5x higher at the centroid at day 2 compared to the centroid at day 3, with a p-value of 0.061 (**Fig 3F**). At the periphery, S10 signal only decreased 1.26x between day 2 and day 3. However, the centroid signal remained significantly higher than the periphery at day 3 (**Fig 3F**). These data suggest that growth is slowing at both spatial locations by day 3 post-inoculation, consistent with our previous results using slowed growth reporters (21, 35), but centroid cells experience a greater decrease in S10 expression between the two experimental timepoints. Despite this global decrease across the microcolony, centroid cells retain more S10 signal across all timepoints relative to peripheral cells.

To explore the relationship between *yopE* and S10 expression within a single microcolony, we generated correlation plots of the mCherry and GFP fluorescent signals at the centroid and periphery. At day 2 post-inoculation, many microcolonies at the centroid displayed high S10 and low *yopE* expression, consistent with our hypothesis (**Fig 3G)**. However, given the smaller sample size of microcolonies within the tissue and heterogeneity between microcolonies at this timepoint, there was not a correlation between signals. At day 3 post-inoculation, more microcolonies are present in the tissue for analysis, and this larger sample size at day 3 allowed us to also analyze cells separately based on levels of *yopE* expression (greater or less than mean fluorescent intensity of 100). *yopE-*expressing cells (MFI ≥100) at the centroid and periphery had a weak positive correlation between *yopE* and S10 levels, with slight positive slopes that appeared quite flat due to low S10 levels **(Fig 3H)**. We noted the S10 levels were higher, and *yopE* levels lower, at the centroid compared to the periphery. In contrast, *yopE* low cells (MFI < 100) had significantly higher levels of S10 expression, as evidenced by a best-fit line with a negative slope **(Fig 3H)**.

These results indicate that two populations emerge within the mouse spleen: bacterial cells with higher *yopE* expression and lower ribosomal activity, and cells with lower *yopE* expression that retain higher levels of ribosomal activity. The second population becomes significantly more pronounced at day 3 compared to day 2, likely due in part to a larger sample size at day 3. We also observed an overall decrease in S10 signals at day 3 **(Fig 3F)** that results in more uniformity across the bacterial populations and made it possible to more clearly distinguish distinct subpopulations of cells.

### T3SS induction slows growth and is associated with decreased S10 expression for the majority of the bacterial population

Outside of a mammalian host, T3SS expression can be induced in *Yersinia* cultures by magnesium oxalate (MgOx) treatment resulting in slowed growth (15, 38). This induction strategy involves adding equimolar amounts of MgCl_2_ and sodium oxalate, which dissociate to form NaCl and MgOx. This strategy will add Na^+^ and Mg^2+^ cations in the absence of added calcium (Ca^2+^) in culture media, resulting in low calcium conditions that are known to trigger high levels of T3SS expression (15, 45). It is important to note that incubation at 37° C induces a low level of T3SS expression, and MgOx addition heightens expression to a growth-inhibited, full induction state. To explore if growth inhibition results in decreased S10 expression, the *yopE::mCherry P_S10_::gfp-ssrA* reporter strain was grown at 37° C in the presence or absence of MgOx to induce T3SS expression. As expected, growth curves demonstrated significant growth inhibition at 3h and 4h post-MgOx treatment (**Fig 4A**). T3SS expression was significantly higher in +MgOx treated samples compared to –MgOx between 2h-4h of treatment, correlating well with growth inhibition **(Fig 4B, 4C)**. However, these data suggest T3SS expression may precede growth inhibition, based on the timing of reporter signal detection (2h) and growth inhibition (3h) **(Fig 4A, 4C)**.

**Fig 4.**
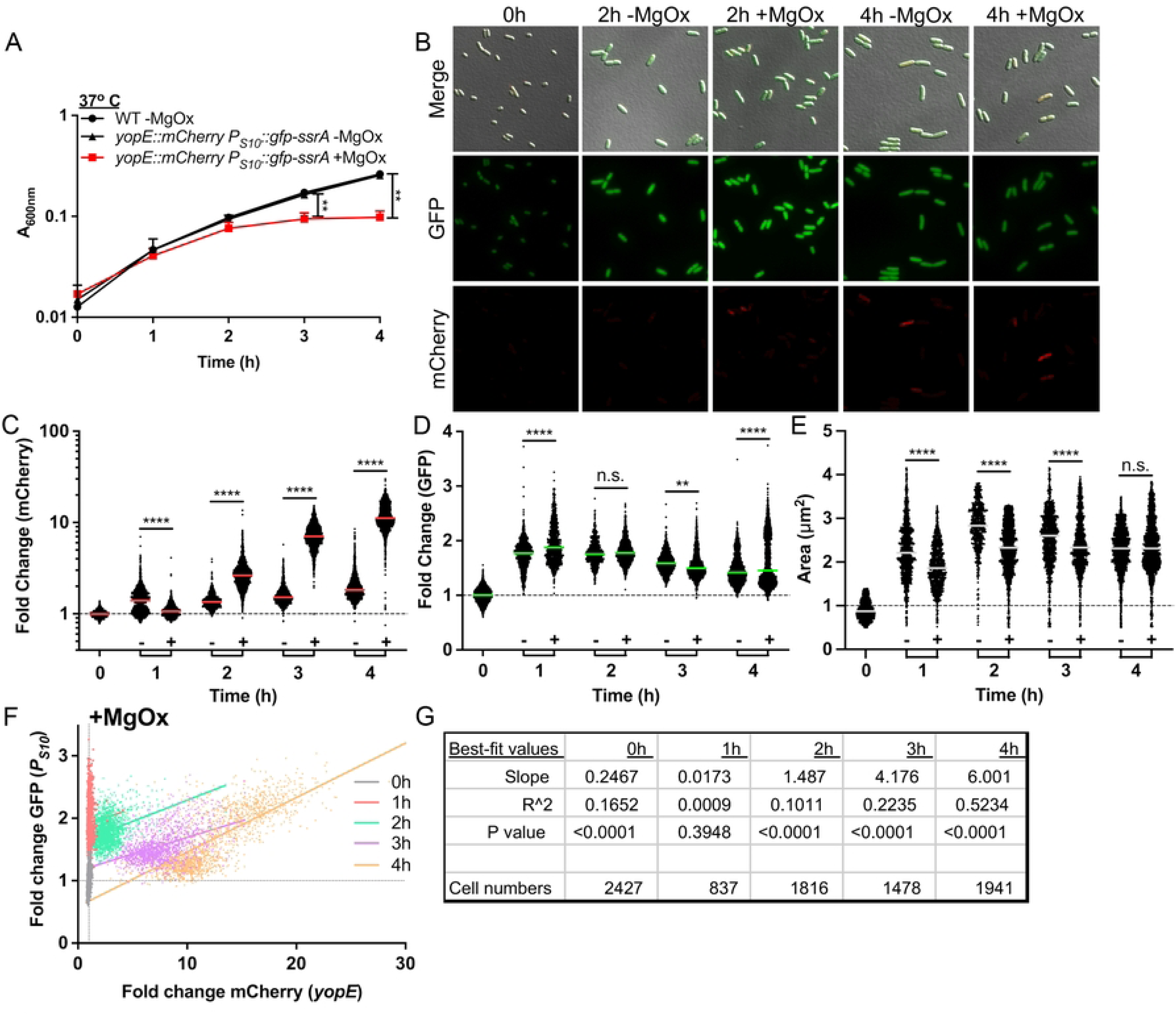
T3SS induction slows growth and is associated with decreased S10 expression for the majority of the bacterial population. WT and *yopE::mCherry P_S10_::gfp-ssrA* strains were cultured at 37° C in the presence (+, T3SS-induced) or absence (-) of MgOx for the indicated timepoints (hours, h). **(A)** Growth curve of strains with and without MgOx. Absorbance (A_600nm_) detected by plate reader at the indicated timepoints. Statistics compare the *yopE::mCherry P_S10_::gfp-ssrA* strain in the presence (+) or absence (-) of MgOx. Mean and standard deviation are shown. **(B)** Representative fluorescence microscopy images of bacteria from **(A)** immobilized on 1% agarose pads. Fold change in **(C)** mCherry (*yopE::mCherry*) reporter signal and **(D)** GFP (*P_S10_::gfp-ssrA*) reporter signal in the absence (-) or presence (+) of MgOx. Values quantified in individual bacteria by fluorescence microscopy. Single cell fluorescence was normalized to the average fluorescent value at 0h (value of 1, represented by dotted line). Each dot: individual cell, horizontal lines: median values. **(E)** Quantification of single cell bacterial areas (µm2) from samples in panels **(C)** and **(D)**. Horizontal lines: median values. **(F)** Correlation plot of fold change in single cell mCherry and GFP fluorescence from bacteria in panels **(C-E**) cultured in the presence (+) of MgOx. **(G)** Linear regression data indicating slope, R2, and significance of the lines of best fit shown in **(F)**. All data represent 3 biological replicates for each strain and condition in this figure. Statistics: **(A)** Two-way ANOVA with Tukey’s multiple comparison test; **(C-E)** Kruskal Wallis one-way ANOVA with Dunn’s post-test; ****p<0.0001, **p<.01, ns: not-significant. **(F,G)** Significantly non-zero slope indicates correlation between values.

Interestingly, while S10 expression generally correlated with growth, it had an inverse relationship with T3SS expression (**Fig 4A, 4D**). At the 1h timepoint when untreated cells exhibited higher median T3SS expression, this population also exhibited lower median S10 expression. As the time course continued, MgOx treated cells increased expression of the T3SS resulting in a concomitant decrease in S10 signal, significantly so by 3h post-treatment when growth was also arrested **(Fig 4A, 4C, 4D)**. Additionally, MgOx treatment resulted in a smaller median cell area, consistent with decreased division rate and decreased metabolic activity (**Fig 4E**) (46). The difference in median cell size was lost at the 4h timepoint, suggesting that untreated cells may also be approaching stationary phase at this timepoint, and are starting to slow their metabolism.

Interestingly, while we observed the median S10 reporter signal was higher in cells treated with MgOx 4h compared to untreated cells, this was largely due to a subset of bacterial cells that retained high S10 expression **(Fig 4D)**. We also noted there were cells present with lower T3SS expression at this timepoint **(Fig 4C**), and we hypothesized this subset with lower T3SS expression may have retained higher translational activity, and thus higher S10 reporter expression. Finally, to explore the relationship between S10 and T3SS expression, we plotted the fold change of each fluorescent marker for individual cells treated with MgOx. In the first hour after treatment S10 expression is strongly induced in all cells, likely capturing their transition out of stationary phase, as seen by a shift up the y-axis (in red) relative to the 0h timepoint (in gray) **(Fig 4F)**. As also seen with transcript levels in **Fig 2D**, S10 expression levels appear highest at 1h-2h of culture (red, green), and gradually begin to decrease **(Fig 4F**). After the initial increase, S10 levels generally decreased as T3SS expression increased; evidenced by a shift in the population towards the bottom right of the graph **(Fig 4F)**.

To our surprise, S10 levels did not decrease universally. At 4h post-treatment (in orange) two populations of cells emerged; a population with low S10 and low T3SS expression and a population with high S10 and high T3SS expression **(Fig 4F)**. In contrast with our hypothesis, this suggests that a subpopulation of cells is capable of expressing both high levels of T3SS and ribosomal protein genes (S10). The linear regression statistics were consistent with a positive correlation between the two reporter signals at the 4h timepoint, based on a significantly non-zero positive slope and R^2^ value of 0.5234 **(Fig 4G)**. There was very little correlation between S10 and T3SS levels in the absence of MgOx based on the slope and R^2^ values of the best-fit lines, likely because T3SS expression was low under these conditions **(S1 Fig)**.

Based on our mouse model datasets, we expected that bacteria grown in T3SS-inducing conditions would exhibit lower S10 reporter activity concomitant with a slowed growth phenotype. Our culture conditions instead indicate that a subset of bacteria exhibit high T3SS activity and high levels of S10 expression. Subpopulations of bacteria with high T3SS expression and high translational activity have been observed in culture (47). However, based on our results, we hypothesize there are kinetic differences in the response and impact of each signal, where prolonged T3SS induction may be necessary to slow the translational activity of an individual cell. We also expect that additional host-derived factors impact the ribosomal content of cells expressing high levels of the T3SS *in vivo*.

### Low levels of T3SS expression are not sufficient to reduce ribosomal reporter activity

We next asked whether the induction of T3SS alone, independent of temperature, is sufficient to impact bacterial growth and S10 reporter expression. Bacteria were incubated in the presence or absence of MgOx at 26° C, absorbance measurements were taken, and cells were collected for fluorescence microscopy. The *yopE::mCherry* strain (lacking *P_S10_::gfp-ssrA*) and non-fluorescent WT strains were used as controls to ensure growth was not impacted by expression of either reporter (*yopE::mCherry* or *P_S10_::gfp-ssrA*). There were no significant differences, indicating the reporters did not impact growth and that MgOx treatment alone, without a temperature shift to 37° C, was not sufficient to inhibit bacterial growth **(S2 Fig, A)**. Fluorescence microscopy indicated MgOx treatment resulted in low levels of *yopE::mCherry* expression **(S2 Fig B, C)**. Levels of T3SS induction were very low relative to MgOx-treated cells at 37° C, and treated cells at 26° C no longer had T3SS induction at 4h post-treatment **(S2 Fig, C)**. In the absence of significant growth inhibition, S10 levels were minimally impacted by treatment, however MgOx-treated cells had significantly lower S10 expression at 3h post-treatment **(S2 Fig, D)**, and we observed a decrease in median cell size 2h-4h after MgOx treatment **(S2 Fig, E)**. These results were somewhat surprising, as we expected addition of MgOx alone would induce significant levels of T3SS expression and subsequent growth inhibition (32, 33). However, since did not observe growth inhibition, it was not surprising that S10 levels remained similar in the presence and absence of MgOx treatment. The decreased median cell size under inducing conditions may also suggest that MgOx treatment, and the release of cations associated with this treatment, has other impacts on cell physiology **(S2 Fig, E)**.

### Loss of the virulence plasmid promotes increased growth rates under T3SS-inducing conditions and is associated with high S10 expression

To assess the role of T3SS expression in growth arrest and subsequent reduction in S10 reporter expression, we transformed the *P_S10_::gfp-ssrA* reporter plasmid into a plasmid-cured *Y. pseudotuberculosis* strain lacking the virulence plasmid (P(-)) (36, 45), a 68kb extrachromosomal plasmid that encodes for the T3SS structural proteins and associated effector proteins (36, 48). It is important to note there are additional genes encoded on the virulence plasmid, notably *yadA*, a temperature-regulated adhesin and known virulence factor that promotes injection of T3SS effector proteins into host cells (31). We initially decided to utilize the P(-) strain for culture-based experiments to assess the role of T3SS expression, as we expected the impact of YadA to be minimal under these conditions.

To confirm that T3SS expression underlies growth arrest, WT *P_S10_::gfp-ssrA* and P(-) *P_S10_::gfp-ssrA* strains were treated with MgOx and grown at 37° C. As expected, the P(-) *P_S10_::gfp-ssrA* strain had significantly higher absorbance values than WT *P_S10_::gfp-ssrA* after 4h culture, indicating the presence of the virulence plasmid significantly impaired growth **(Fig 5A)**. To determine if differences in growth rates correlated with changes in S10 levels, we detected S10 reporter expression by fluorescence microscopy. The WT *P_S10_::gfp-ssrA* and P(-) *P_S10_::gfp-ssrA* strains had similar levels of S10 expression at 0h, but S10 expression levels in the P(-) strain increased at 2h, and were significantly higher than WT at 4h growth **(Fig 5B)**. This was consistent with the observed growth inhibition at 4h treatment of the WT strain **(Fig 5A)**, again indicating that decreased S10 reporter expression correlated with growth arrest, and that heightened levels of S10 expression are detected in faster-growing cells. Similar to results in **Fig 3**, we also observed a great deal of heterogeneity within the bacterial populations. While the overall median value of S10 reporter expression was higher in P(-) cells at 4h, there was a subpopulation of cells with low S10 levels as well **(Fig 5B)**. This could suggest there are both fast and slow growing cells present within the population. Collectively, these data indicate that the growth inhibition associated with expression of virulence plasmid-encoded genes correlates with a reduction in S10 levels.

**Fig 5.**
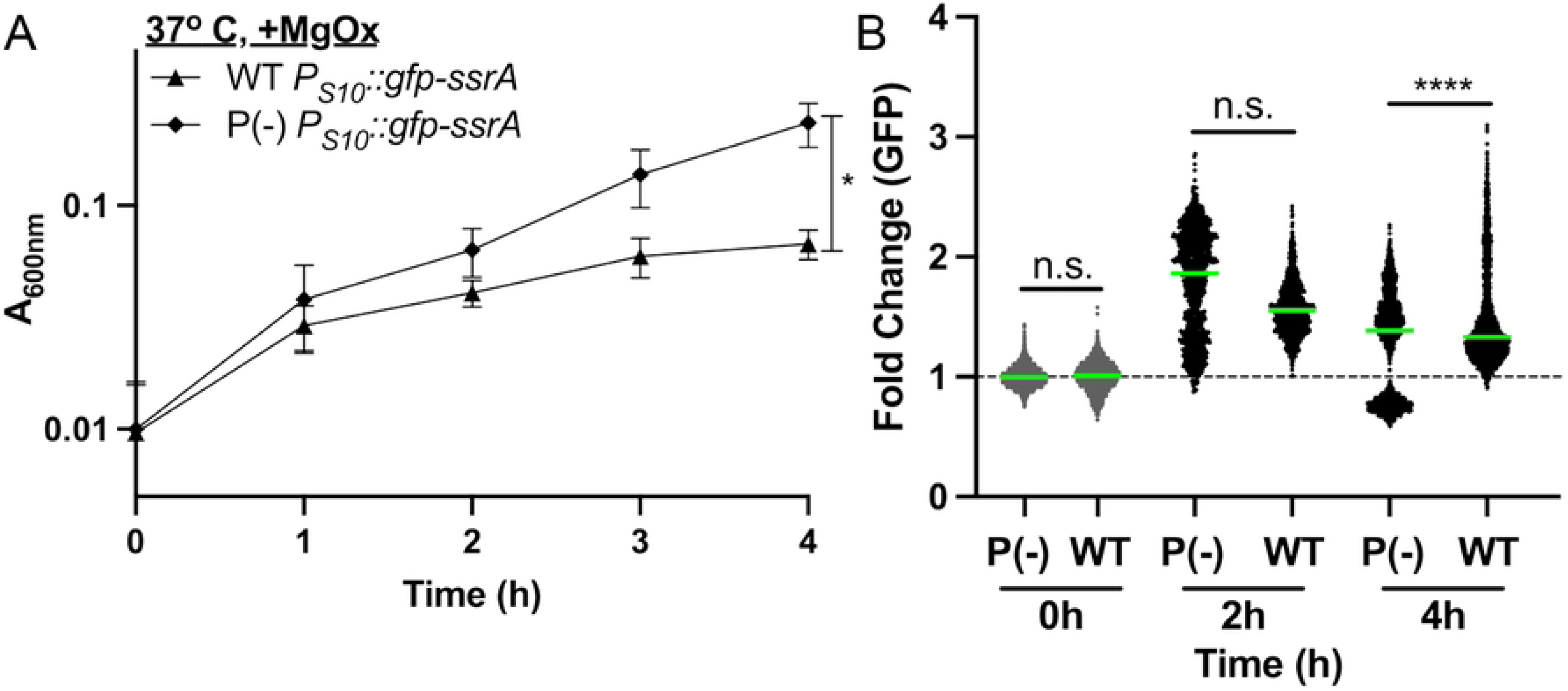
Loss of the virulence plasmid is associated with increased growth rates and heightened S10 expression under T3SS-inducing conditions. WT, *P_S10_::gfp-ssrA,* and P(-) *P_S10_::gfp-ssrA* (virulence plasmid-cured) strains were cultured at 37° C in the presence (+) of MgOx for the indicated timepoints (hours, h). **(A)** Growth curve of strains, absorbance (A_600nm_) was detected by plate reader at the indicated timepoints. Statistics compare the P(-) strain to other strains. Mean and standard deviation are shown. **(B)** Fluorescence microscopy to quantify single cell fold change in GFP (*P_S10_::gfp-ssrA*) reporter signal. Single cell fluorescence was normalized to the average fluorescent value of the WT strain at 0h (value of 1, represented by dotted line). Each dot: individual cell, horizontal lines: median values. All data represent 3 biological replicates for each strain and timepoint in this figure. Statistics: **(A)** Two-way ANOVA with Tukey’s multiple comparison test; **(B)** Kruskal Wallis one-way ANOVA with Dunn’s post-test; ****p<0.0001, *p<.05, ns: not-significant.

### T3SS induction reduces gentamicin susceptibility

We hypothesized that the growth arrest observed with MgOx treatment may be sufficient to decrease antibiotic susceptibility, based on known associations between slowed growth, reduced metabolic activity, and decreased antibiotic susceptibility (49, 50). To test this hypothesis, we cultured the *yopE::mCherry P_S10_::gfp-ssrA* strain in the presence and absence of MgOx, then exposed bacteria to either gentamicin or ciprofloxacin **(Fig 6A)**. These antibiotics were chosen because they are bactericidal, albeit with distinct drug targets, the ribosome and DNA gyrase, respectively. MgOx treatment significantly reduced the susceptibility of *Y. pseudotuberculosis* to gentamicin, as evidenced by increased survival during gentamicin exposure **(Fig 6B)**. MgOx treatment was also sufficient to promote tolerance to ciprofloxacin **(Fig 6C)**. However, it has been well-established that the presence of metal cations, including Mg^2+^, can reduce ciprofloxacin activity (51). To determine whether altered antibiotic susceptibility was linked to T3SS expression or instead impacted by divalent cations, we performed additional experiments with a Δ*lcrF* strain, which lacks T3SS induction (52). The Δ*lcrF* strain had similar growth kinetics in the presence and absence of MgOx, displayed increased sensitivity to gentamicin relative to the WT strain, and Δ*lcrF* gentamicin sensitivity was similar in the presence or absence of MgOx **(Fig 6D)**, together indicating that the reduced sensitivity of the WT strain to gentamicin was linked to T3SS induction. In contrast, Δ*lcrF* had similar ciprofloxacin sensitivity compared to the WT strain in the absence of MgOx, and in the presence of MgOx, the Δ*lcrF* continued to replicate during ciprofloxacin exposure, consistent with resistance **(Fig 6E)**. This indicates that reduced ciprofloxacin sensitivity was instead due to cation interaction with the antibiotic, and not T3SS expression.

**Fig 6.**
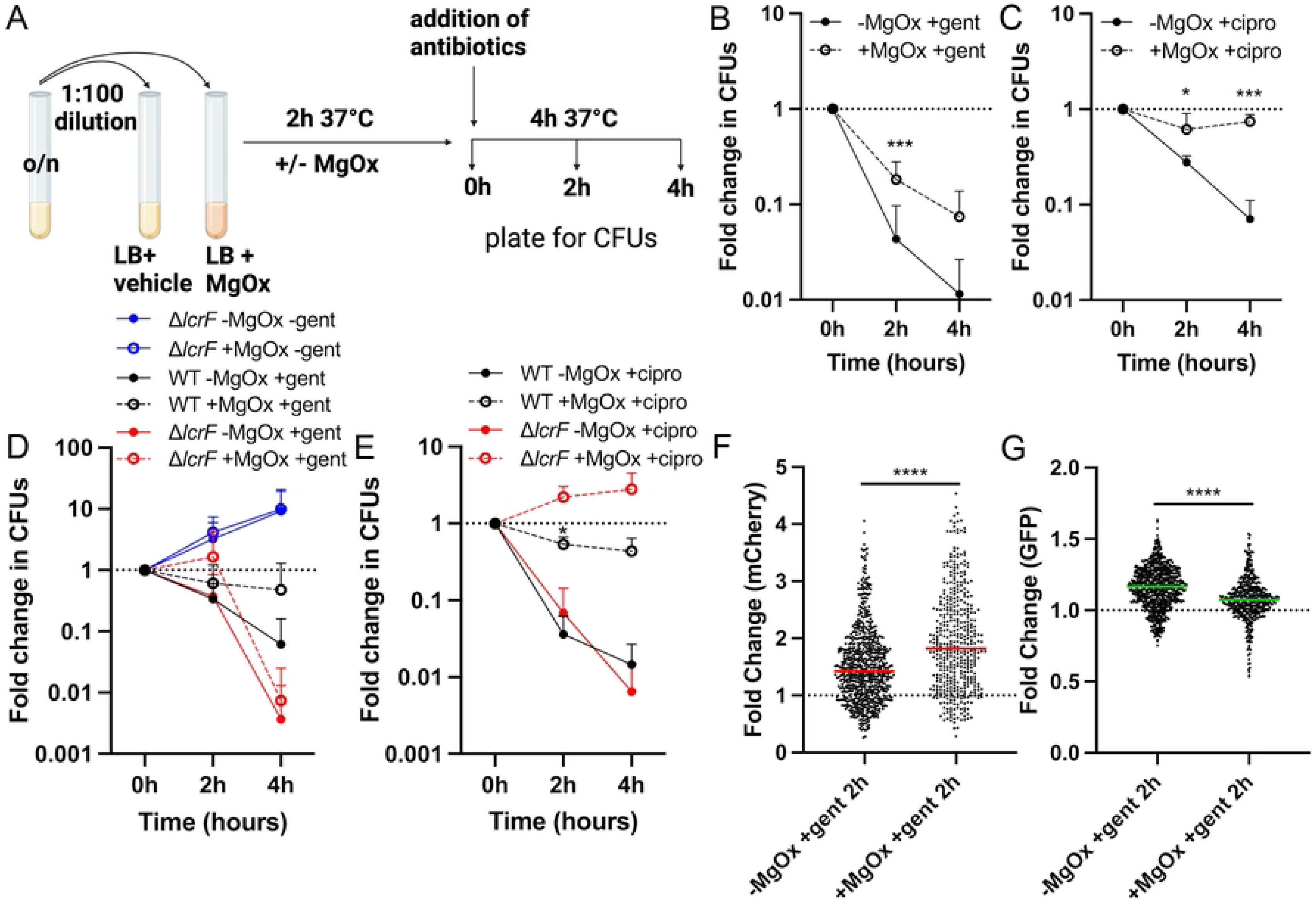
T3SS induction reduces gentamicin susceptibility. Strains were grown in the presence (+) or absence (-) of MgOx for 2h at 37° C. Antibiotics were added (gentamicin: gent, ciprofloxacin: cipro) and strains were cultured an additional 4h. Samples were taken at 0h (addition of antibiotics), 2h, and 4h to quantify CFUs or quantify fluorescence by microscopy. Fold change in CFUs is calculated by normalizing the number of viable bacteria at 2h and 4h to the number of viable bacteria at 0h. **(A)** Experimental design. **(B)** *yopE::mCherry P_S10_::gfp-ssrA* strain +/- MgOx, gent treatment. **(C)** *yopE::mCherry P_S10_::gfp-ssrA* strain +/- MgOx, cipro treatment. **(D)** Comparison between non-fluorescent WT and Δ*lcrF* strains, +/- MgOx, gent treatment. **(E)** Comparison between non-fluorescent WT and Δ*lcrF* strains, +/- MgOx, cipro treatment. Fold change in **(F)** mCherry (*yopE*) and **(G)** GFP (S10) signal from cells represented in **(B)** was calculated by normalizing the single cell fluorescence to the average fluorescent value at 0h (value of 1, represented by dotted line). Each dot: individual cell, horizontal lines: median values. Dead cells were excluded using the fixable blue dead cell stain. Statistics: **(B-E)** Two-way ANOVA with Tukey’s multiple comparison test, statistics shown represent comparisons +/- MgOx; **(F, G)** Mann-Whitney; ****p<0.0001, ***p<.001, *p<.05.

To determine if gentamicin treatment selected for bacterial survivors with heightened levels of T3SS and lower levels of S10, we performed fluorescence microscopy with the *yopE::mCherry P_S10_::gfp-ssrA* strain after 2h of gentamicin treatment. 2h was chosen because we observed significant changes in viability (**Fig 6B**) and because this timepoint correlates with the 4h MgOx timepoint of **Fig 4 and Fig 5**. We quantified fluorescent reporter expression of surviving, viable bacterial cells by incorporating a live/dead fluorescent dye to exclude non-viable cells from our analyses. Bacterial cells that survived gentamicin treatment had significantly higher mCherry **(Fig 6E)** and lower S10 **(Fig 6F)** reporter signal in the MgOx treatment group compared to untreated, consistent with increased survival of slower-growing cells with high T3SS and lower S10.

### Droplet-based microfluidics can be used to model clustered bacterial growth

One limitation of *in vivo* infection models is the inability to track growth and reporter expression profiles over time. In contrast, culture-based experiments can enable kinetic analyses, but lack the clustered spatial architecture of bacteria seen within the host environment. To circumvent these issues, droplet-based microfluidics can be used to generate local agar environments that force bacteria to grow in tightly-clustered colonies, as they would within host tissues (20, 53). Here, we employed microfluidic droplets to observe colony growth and reporter expression over time during incubation at 26° C and 37° C, and at 37° C in the presence and absence of MgOx. Agarose-based microfluidics droplets were generated as previously described, where liquid and oil phases intersect to generate oil-encapsulated spherical droplets, and an appropriate bacterial density in the liquid phase was used to ensure a majority of droplets were seeded with a single bacterial cell (53). Oil was removed from the exterior of droplets to allow free diffusion of molecules into and out of the droplets during culture (53). We hypothesized that MgOx addition during culture at 37° C would result in high levels of T3SS induction, growth inhibition, and lower S10 levels. We further reasoned that increased accumulation of MgOx might promote heightened T3SS activity at the periphery of microcolonies. Based on results in culture, we hypothesized that the temperature shift to 37° C alone would not significantly delay microcolony growth relative to 26° C, since this should result in intermediate T3SS induction.

We first explored if temperature impacted microcolony growth within droplets by encapsulating the *yopE::mCherry P_S10_::gfp-ssrA* strain into droplets and culturing at 26° C or 37° C. Cultures were sampled every two hours to quantify the microcolony area within droplets, T3SS reporter signal, and S10 reporter signal by fluorescence microscopy. Microcolonies grew at both temperatures, but were significantly larger at 37° C (**Fig 7A**). While microcolonies had significant T3SS induction at 4h and 6h of growth at 37° C (**Fig 7B**), there was no change in S10 reporter signal between conditions (**Fig 7C**). These results corroborate planktonic cell culture experiments where growth at 37° C moderately induces T3SS expression but does not reduce growth rate **(Fig 4A-C**).

**Fig 7.**
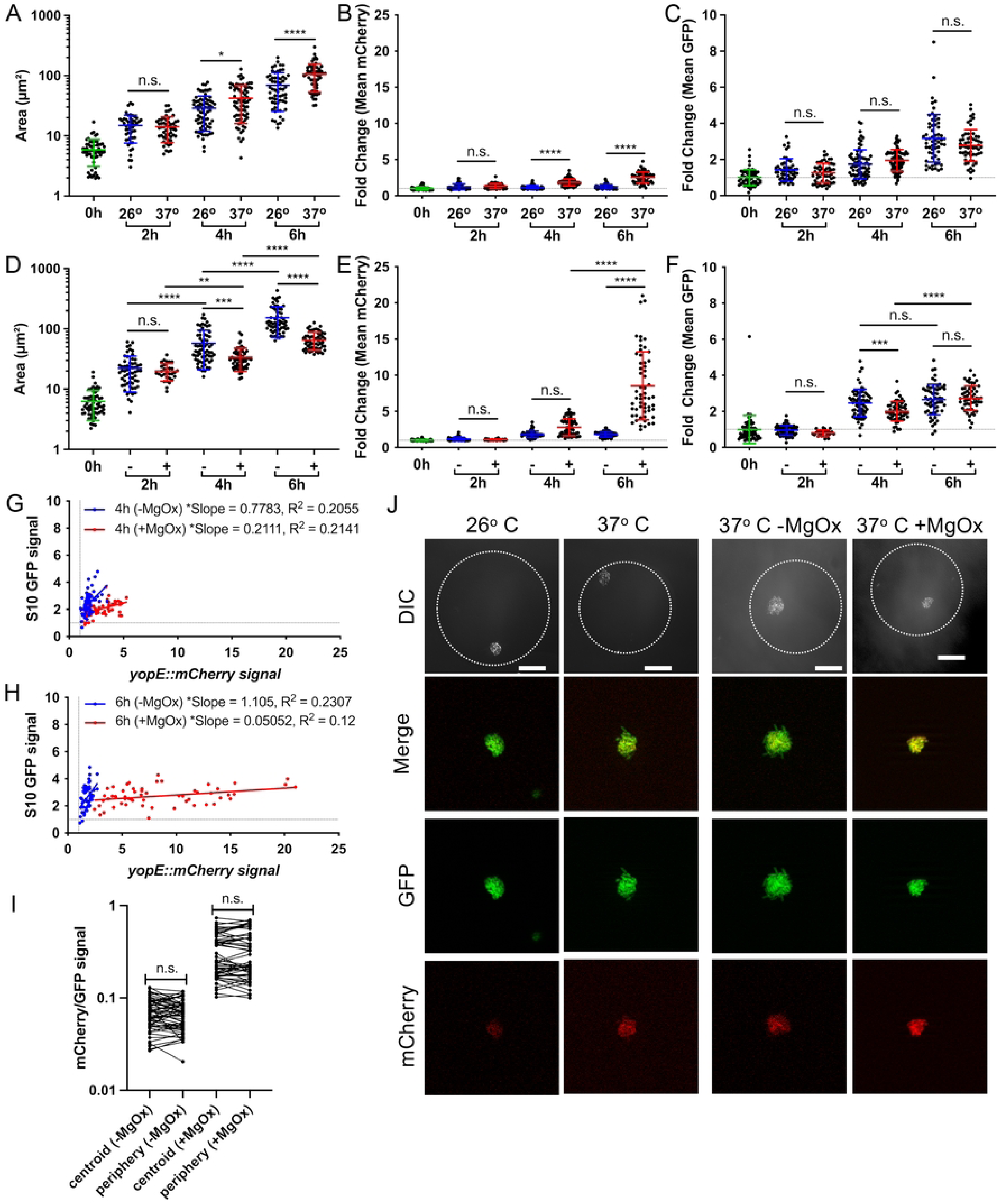
Droplet-based microfluidics can be used to model clustered bacterial growth. The *yopE::mCherry P_S10_::gfp-ssrA* strain was encapsulated into droplets and cultured for at the indicated temperatures in the presence (+) or absence (-) of MgOx. At the indicated times aliquots were taken and bacterial growth and reporter expression were quantified by fluorescence microscopy. Growth at 26° C was compared to 37° C, and **(A)** area (um2), **(B)** fold change in mCherry signal (relative to 0h), and **(C)** fold change in GFP signal (relative to 0h) were quantified. Horizontal dotted lines represent a value of 1, the initial average value at time 0h. Growth at 37° C in the presence (+) or absence (-) of MgOx was compared, and **(D)** area (um2), **(E)** fold change in mCherry signal (relative to 0h), and **(F)** fold change in GFP signal (relative to 0h) were quantified. **(G, H)** Linear regression data indicating slope and R2 value of best fit lines for GFP compared to mCherry signal of microcolonies grown **(G)** 4h or **(H)** 6h at 37° C, (+) or (-) MgOx. **(I)** mCherry/GFP signal ratios are shown at the indicated spatial locations for microcolonies grown -/+ MgOx. **(J)** Representative images of microcolonies grown within droplets under the indicated conditions, all images represent 6h timepoints. Differential Interference Contrast (DIC) shown alongside GFP, mCherry, and merged fluorescent channels. Dotted line outlines the periphery of each droplet. Scale bar: 20um. All data represent 2 biological replicates for each condition in this figure. Mean and standard deviation are shown. Each dot represents an individual microcolony. Statistics: **(A-F)** Two-way ANOVA with Tukey’s multiple comparison test; **(G, H)** Linear regression, significantly non-zero slope indicates correlation between values; **(I)** Wilcoxon matched-pairs; ****p<0.0001, ***p<.001, **p<.01, *p<.05, ns: not-significant.

While temperature shift would be sensed by the entire population during infection, local chemical mediators, such as those released from surrounding neutrophils or monocytes (20), can create chemical gradients experienced most strongly by peripheral cells in the microcolony. To determine if agarose droplets limit soluble molecule diffusion and generate similar chemical gradients, MgOx was added to droplet cultures at 37° C. We observed significant microcolony growth inhibition at 4h and 6h post MgOx treatment (**Fig 7D**) and high levels of T3SS induction (**Fig 7E**) after 6h of culture. Again, these results match planktonic cell culture experiments where MgOx treatment at 37° C strongly induces T3SS expression and results in growth inhibition (**Fig 4A-C**). Surprisingly, S10 expression did not correlate with growth inhibition at all timepoints. At the 4hr timepoint S10 expression was significantly lower in the growth inhibited +MgOx group, but this difference disappeared after 6 hours of culture, due to increased expression between timepoints in the treated group (**Fig 7F**). These results suggest that the MgOx-treated group may have experienced a “second wind” and resumed growth between the timepoints. Interestingly, correlation plots comparing T3SS and S10 expression suggested droplets represented the phenotypes observed within the mouse spleen better than culture, with detection of T3SS low, S10 high cell populations under uninduced conditions (blue) (**Fig 7G, 7H**).

These data were analyzed using the mean fluorescent intensity of each fluorescent signal across an individual microcolony. To explore how local position in the growing microcolony impacts reporter expression, we analyzed individual droplet microcolonies by quantifying their centroid and peripheral fluorescence values after 6h of culture, when T3SS reporter signal was highest. While we have consistently seen a significant increase in T3SS signal at the periphery of microcolonies within host tissues (**Fig 3C, 3D, 3E**) (20), we did not observe any spatial difference in mCherry/GFP (T3SS/S10) reporter signal within droplets during growth at 37° C in the presence or absence of MgOx (**Fig 7I, 7J**). This suggests that MgOx diffuses readily through droplets, and does not accumulate in a particular spatial location, which is consistent with its size relative to the presumed porosity of the agarose droplet particles. This could be beneficial for future applications of this technology, where researchers could generate uniform microenvironments within a single droplet or even droplet configurations. These results also suggest that contact with the agarose matrix of droplets, which could mimic surface contact, is not sufficient to mimic neutrophil contact within host tissues. Modifications to the droplet system would be needed to model direct host cell contact.

## Discussion

In this study, we sought to understand the impact of T3SS-induced growth arrest on levels of ribosomal protein transcripts and determine whether reduced expression of ribosomal protein genes correlated with reduced antibiotic susceptibility. We generated a fluorescent transcriptional reporter to detect levels of expression of the S10 ribosomal protein operon and found high levels of T3SS induction and expression were required for growth arrest and reduced expression of S10. We observed heterogeneity in single cell gene expression patterns across the bacterial population in the mouse spleen, in an agarose droplet model of clustered bacterial growth, and during planktonic growth in culture. Importantly, we show high levels of expression of the T3SS are sufficient for growth arrest, expression of the T3SS promotes antibiotic tolerance, and bacterial cells that survive antibiotic treatment have heightened levels of T3SS and lower levels of S10 expression.

The reporter construct generated here was used to mark cells that had a rapid-growth phenotype compared to slower-growing bacterial cells. While this construct utilized a promoter upstream of the *rpsJ*/S10 gene, it remains unclear if this reporter detects altered ribosome levels. A previous study by Manina et al. used a similar approach, but instead generated a fluorescent transcriptional reporter for ribosomal RNA in *Mycobacterium tuberculosis*, which may better indicate overall ribosome numbers since rRNA levels typically dictate ribosome biogenesis rates (43). Interestingly, in studies focused on regulation of ribosomal components, it was shown that the *rpsJ*/S10 promoter has unique regulation relative to other ribosomal proteins, and can instead mirror transcript levels of rRNA due to DksA- and ppGpp-dependent transcriptional regulation of *rpsJ* expression (40, 54). The ppGpp-dependent regulatory element found in the leader sequence is included in our reporter, but it is important to note that the first ten amino acids of RpsJ were not included. These amino acids form a hairpin structure that is important for translational regulation (54), thus our reporter is indicating transcriptional activity.

Similar to our results with *Y. pseudotuberculosis*, populations of *M. tuberculosis* exhibit a great deal of heterogeneity in expression of ribosomal genes, which can be amplified under stressful conditions, including infection of the mouse lung and stationary phase culture (43). Following administration of isoniazid in culture, the majority of surviving bacteria expressed higher levels of rRNA, despite an enrichment in the amount of nongrowing cells (43). We also detected heterogeneity within the surviving bacterial population following gentamicin treatment, but overall found increased T3SS and decreased S10 expression in surviving cells, which may better align with results in *Salmonella* showing metabolically-active cells with decreased growth rates have the highest survival rates during antibiotic exposure (55–57).

Previous studies utilizing *Salmonella* have shown that T3SS expression is sufficient to decrease bacterial growth rates, indicating T3SS expression can come at a fitness cost (1, 58, 59). Similar results have been shown with *Yersinia* in culture, where bacterial cells injecting T3SS effectors are growth arrested (14). Here we sought to determine if T3SS-induced growth arrest impacted ribosomal protein expression, since we detected reduced transcript levels of ribosomal protein genes in slower growing, stationary phase *Y. pseudotuberculosis* cultures. Additional studies are needed to compare stationary phase and T3SS-induced growth arrest, but based on our experiments, we predict T3SS induction generates a pronounced, distinct, expression profile with specific effects on cell division machinery and metabolism. It will also be important to separate effects of cations generated during T3SS induction from effects of expression of the T3SS itself, which could be uncoupled using the Δ*lcrF* deletion strain described here (52).

In *Salmonella* systems, it has also been shown that T3SS expression is linked to decreased antibiotic susceptibility (1, 2, 60). In *Yersinia*, much is known about regulation of the T3SS at the molecular level, and it is well appreciated that dynamic expression of this system is required for virulence (22, 61, 62), and so regulation of the system was not a focus of this study. Instead we sought to link altered T3SS expression levels with differences in S10 reporter signal as a measure of growth rate, and determine whether this impacts changes in antibiotic susceptibility. Further, we chose to focus on a ribosome-targeting antibiotic, gentamicin, to determine if distinct levels of ribosomal proteins resulted in differential antibiotic susceptibility. Our finding that gentamicin treatment resulted in survival of cells with lower levels of S10 reporter expression is consistent with the model that cells with lower levels of ribosomes, and lower level of antibiotic target in this case, were better able to survive antibiotic treatment.

Although gentamicin sensitivity can be linked to differences in drug uptake, it has also been recently shown that susceptibility can also depend on ribosomal protein levels, consistent with our results (63). Based on our *in vitro* characterization, our reporter is marking rapidly-dividing cells as intended, but additional experiments would be needed to determine how well this correlates with ribosome numbers.

## Materials and Methods

### Bacterial strains & growth conditions

The WT *Y. pseudotuberculosis* strain, IP2666, was used throughout (20, 64). For *in vitro* experiments with fluorescent reporters, bacteria were grown in LB broth overnight (16h) at 26° C with rotation, sub-cultured 1:100, and then incubated at either 26° C or 37° C with rotation in the absence (-MgOx) or presence of magnesium oxalate (+MgOx: 20mM MgCl_2_ and 20mM sodium oxalate (15, 33)) to induce T3SS expression for the indicated timepoints. Equivalent volume (200µl) dH_2_O was added to -MgOx tubes. Bacterial growth was assessed based on absorbance at 600nm using a Synergy H1 microplate reader (Biotek). For mouse infection experiments, bacteria were grown overnight (16h) to post-exponential phase in 2xYT broth (LB, with 2x yeast extract and tryptone) at 26° C with rotation as previously described (20). The *P_S10_::gfp-ssrA* reporter was expressed from the low copy pMMB67EH plasmid; all cultures with strains containing this construct were supplemented with carbenicillin (100ug/ml).

### RNA isolation and qRT-PCR

Bacterial cells were grown in LB broth for the indicated time points, pelleted, resuspended in Buffer RLT (QIAGEN) + ß-mercaptoethanol, and RNA was isolated using the RNeasy kit (QIAGEN). DNA contamination was eliminated using the on-column DNase digestion kit (RNase-free DNase set, QIAGEN). RNA was reverse transcribed using the Protoscript II First Strand cDNA synthesis kit (NEB). qRT-PCR reactions were prepared with 0.5 µM of forward and reverse primers added to cDNA samples and amplification using SYBR Green (Applied Biosystems). *rpoC* was used as an endogenous control since 16S transcript levels were detected as a target gene. Control samples were prepared that lacked reverse transcriptase to confirm DNA was eliminated from samples. Reactions were carried out using the StepOnePlus Real-Time PCR system (Applied Biosystems), and relative comparisons were obtained using the ΔΔC_T_ or 2^-ΔCt^ method. Kits were used according to manufacturers’ protocols.

### RNA-seq

Overnight cultures (16h, stationary phase) were sub-cultured 1:100 into fresh LB and grown an additional 4h at 26° C with rotation to generate exponential phase samples. Approximately 10^5^ bacterial cells were collected from cultures and RNA was isolated as described above. Purified RNA was shipped to Novogene (Novogene Corporation Inc), for library preparation, sequencing, and bioinformatic analysis. Sequencing reads were aligned to IP2666 genome assembly GCA_003814345.1. The DESeq2 method of pairwise comparisons between treatment groups (65) was used to determine significant differences in transcript levels between stationary and exponential phase cells. An adjusted p-value of 0.05 was considered significant, and hits were determined based on ≥2 fold change, and transcripts with less than 100 mean reads under both conditions were excluded. Full datasets are in **S1 Table and S2 Table.** KEGG pathway analyses were utilized to determine the biological pathways overrepresented in exponential phase compared to stationary phase.

### Generation of reporter strains

The following *Y. pseudotuberculosis* strains were previously described: the strain lacking the virulence plasmid (P(-)) (66), the *yopE::mCherry* strain (20, 24), and the Δ*lcr* strain (52). The *P_S10_::gfp-ssrA* reporter was constructed by fusing the *P_S10_*promoter to *gfp-ssrA* by overlap extension PCR and ligating this fragment into the multiple cloning site of the low copy pMMB67EH plasmid. *P_S10_::gfp-ssrA* strains were constructed by transformation of this plasmid into *Y. pseudotuberculosis* strains using a previously described protocol (67). The constitutive GFP construct in pACYC184 was used as a template for mutagenesis to generate the *gfp-ssrA* construct (20), which was constructed by inserting a ssrA-tag at the 3’ end of *gfp* between the last coding amino acid and the stop codon (21).

### Fluorescence microscopy: bacteria

Bacteria were pelleted, resuspended in 4% paraformaldehyde (PFA) and incubated overnight at 4° C for fixation. PFA was removed and bacteria were resuspended in PBS prior to imaging. Agarose pads were prepared to immobilize bacteria for imaging by solidifying a thin layer of 25µl 1% agarose in PBS between a microscope slide and coverslip (68). Once solidified, coverslips were removed, bacteria were added, coverslips were replaced, and bacteria were imaged with a 63x oil immersion objective, using a Zeiss Axio Observer 7 (Zeiss) inverted fluorescent microscope with XCite 120 LED boost system and an Axiocam 702 mono camera (Zeiss). Volocity image analysis software was used to quantify the fluorescent signal associated with individual bacterial cells (68). WT non-fluorescent cells were grown in parallel for all experiments to detect baseline background fluorescence.

### Antibiotic susceptibility experiments

Overnight cultures were diluted 1:100 into fresh LB treated with MgOx (20mM MgCl + 20mM NaOx) or the same volume of H_2_O (-MgOx) and grown at 37°C to induce the T3SS. After 2h antibiotics were added (gentamicin: 10µg/mL (MBC_99_), ciprofloxacin: 0.1µg/mL (MBC_90_)); this represents the 0h timepoint. Samples were taken at 0h, 2h, and 4h to plate for CFUs and for staining with the Live/Dead Fixable Blue Dead Cell Stain Kit (Invitrogen). After washing in PBS, bacterial samples were stained for 30 minutes, protected from light, and washed in PBS before fixation in 4% PFA. Fixed samples were imaged as described above on agarose pads, and cells with detectable blue fluorescence (non-viable) were excluded from single cell analyses.

### Murine model of systemic infection

Six to eight-week old female C57BL/6 mice were obtained from Jackson Laboratories (Bar Harbor, ME). All animal studies were approved by the Institutional Animal Care and Use Committee of Johns Hopkins University. Mice were injected intravenously into the tail vein with 10^3^ bacteria for all experiments. At the indicated timepoints post-inoculation (p.i.), spleens were removed. ½ of each spleen was homogenized and plated to determine CFU/spleen. Intact tissue (remaining ½ spleen) was processed for fluorescence microscopy as described below.

### Fluorescence microscopy: host tissues

Spleens were harvested and immediately fixed in 4% PFA by incubating overnight at 4° C. Tissues were frozen-embedded in Tissue Plus O.C.T. compound (Fisher Scientific) and cut by cryostat microtome into 10µm sections. Two sections were analyzed for each spleen. To visualize reporters, sections were thawed in PBS at room temperature, stained with Hoechst at a 1:10,000 dilution to detect host cell nuclei, washed in PBS, and coverslips were mounted using ProLong Gold (Life Technologies). Tissue was imaged as described above, with an Apotome.2 (Zeiss) for optical sectioning.

### Image analysis

Volocity image analysis software was used to quantify microcolony areas and fluorescence as previously described (3, 20). Two cross-sections were analyzed for each spleen, and every bacterial cell or clustered microcolony was imaged and analyzed within these sections. Spatial analyses were completed using Volocity software by sampling 4, ∼4µm^2^ regions of interest (ROI) at both the centroid and periphery of each microcolony, and calculating the ratio of mCherry/GFP signal intensity in each ROI. Centroid measurements were captured in a cluster of 4 ROIs around the geometric centroid of each microcolony. Peripheral measurements were taken approximately every 90° around each microcolony, sampling bacterial cells in contact with host cells. Each mCherry/GFP ratio was averaged to generate an average peripheral or centroid measurement for each microcolony, and these values were plotted. Individual fluorescent channel or spatial location values were also plotted using these ROIs. Fluorescent signals were quantified the same way from microcolonies within agarose droplets.

### Droplet generation

CAD-designed microfluidics chips with 38 independent devices controlled by two input ports were constructed according to published protocols (69, 70) at the FabLab Micro and Nano fabrication laboratory at the University of Maryland, and devices were used as previously described (53). Prior to droplet generation, each device was primed with HFE-7500 Novec oil (Oakwood Chemical) by connecting a 1ml syringe fitted with a 26-gauge needle to each input port via polyethylene (PE 20) tubing (BD). A third tube was inserted into the droplet collection port and waste was collected in a 1.5 ml microcentrifuge tube. After priming, the syringe connected to the oil-phase port was replaced with a syringe containing 1.5% FluoroSurfactant in HFE7500 Novec oil by removing the tubing from the needle of the priming syringe and placing the tubing onto the 1.5% FluoroSurfactant syringe. The syringe connected to the aqueous phase port was replaced with a 1 ml syringe containing bacteria diluted 1:2000 in 1% ultra-low-melt agarose, yielding approximately 5 x 10^6^ cells/ml. The oil-phase and aqueous-phase syringes were loaded onto separate programmable syringe pumps (Harvard Apparatus 11 Elite Series) set to 700 µl/hour and 200 µl/hour respectively. A heat lamp (VWR) was placed at a short distance (∼ 0.5 m) from the syringe pump and tubing to prevent premature solidification of the agarose. Droplets were collected in a 1.5 ml microcentrifuge tube and incubated at 4° C for 30 minutes to solidify.

### Oil removal

Oil was removed from the droplets as previously described (53). To remove oil from 200 µl of droplets, aliquots of 100 µl of droplets were moved into two separate 1.5 ml microcentrifuge tubes and 500 µl of 10% 1H, 1H, 2H, 2H-Perfluoro-1-octanol (PF) (Sigma) in Novec 7500 oil was added to each. The tubes were shaken vigorously and centrifuged for 30 seconds at 250 RCF. 400 µl PBS was added and the droplets were flicked into suspension then centrifuged for 30 seconds at 250 RCF. The PBS/droplet layers from each tube were combined in a new 1.5 ml microcentrifuge tube, centrifuged for 30 seconds at 250 RCF, and washed with 1 ml PBS. The remaining droplets were resuspended in 700 µl 2xYT broth.

### Magnesium oxalate droplet experiments

∼400 µl of droplets were generated and separated as 200 µl aliquots resuspended in 1.5 ml microcentrifuge tubes containing 700 µl 2xYT. Both tubes were shaken gently then rotated at 37° C. At 2 hours, MgCl_2_ (20mM) and sodium oxalate (20mM) were added to one tube while an equivalent amount of ddH_2_O (200µl) was added to the untreated tube. Samples were taken at 2-hour intervals by removing the tubes from the incubator, aliquoting 100 µl of each 2xYT-droplet mixture into new microcentrifuge tubes, centrifuging the tubes for 30 seconds at 250 RCF, and resuspending the droplets in 100 µl of 4% PFA.

## Author Contributions

Conceptualization: JG, KMD; Formal Analysis: JG, RAS, KMD; Funding Acquisition and Supervision: KMD; Investigation: JG, RAS, KLC, REB, KMD; Methodology: JG, RAS, RCH, SC, AAC, JMS, KMD; Project Administration: KMD; Visualization: JG, RAS, KLC, KMD; Writing – Original Draft Preparation: JG, RAS, KMD; Writing – Review & Editing: JG, RAS, KLC, REB, JMS, WEB, RG, KMD.

## Acknowledgments

We thank the members of the Davis lab, who provided feedback and suggestions throughout the steps of manuscript preparation. We would also like to thank members of the Pekosz, Klein, and Baumgarth labs for valuable feedback during manuscript preparation, and thank Dr. Patrick Kreisher in particular for his insight into cations. Thank you to Dr. Vicki Auerbuch Stone and Dr. Joan Mecsas for providing Δ*lcr* mutant strains. The authors of this manuscript declare no conflicts of interest.

## Supporting Information Captions

**S1 Table. Full RNAseq dataset: list of genes with heightened reads in exponential phase. S2 Table. Full RNAseq dataset: list of genes with heightened reads in stationary phase.**

**S1 Fig. S10 and T3SS reporter signals do not correlate in the absence of MgOx treatment. A)** Correlation plot of fold change in single cell mCherry and GFP fluorescence from bacteria cultured in the absence (-) of MgOx. **(B)** Linear regression data indicating slope, R2, and significance of the lines of best fit shown in **(A).** Data represents 3 biological replicates for each strain and condition. Significantly non-zero slope indicates correlation between values.

**S2 Fig. Low levels of T3SS expression were not sufficient to reduce ribosomal reporter activity.** WT, *yopE::mCherry,* and *yopE::mCherry P_S10_::gfp-ssrA* strains were cultured at 26° C in the presence (+) or absence (-) of MgOx for the indicated timepoints (hours, h). **(A)** Growth curve of strains with and without MgOx. Absorbance (A_600nm_) detected by plate reader at the indicated timepoints. Statistics compare the *yopE::mCherry P_S10_::gfp-ssrA* strain in the presence (+) or absence (-) of MgOx. Mean and standard deviation are shown. **(B)** Representative fluorescence microscopy images of bacteria from **(A)** immobilized on 1% agarose pads. Fold change in **(C)** mCherry (*yopE::mCherry*) reporter signal and **(D)** GFP (*P_S10_::gfp-ssrA*) reporter signal in the absence (-) or presence (+) of MgOx. Values quantified in individual bacteria by fluorescence microscopy. Single cell fluorescence was normalized to the average fluorescent value at 0h (value of 1, represented by dotted line). Each dot: individual cell, horizontal lines: median values. **(E)** Quantification of single cell bacterial areas (µm2) from samples in panels **(C)** and **(D)**. Horizontal lines: median values. All data represent 3 biological replicates for each strain and condition in this figure. Statistics: **(A)** Two-way ANOVA with Tukey’s multiple comparison test; **(C-E)** Kruskal Wallis one-way ANOVA with Dunn’s post-test; ****p<0.0001, ***p<.001, **p<.01, ns: not-significant.

